# Phage-antibiotic combinations to control *Pseudomonas aeruginosa-Candida* two-species biofilms

**DOI:** 10.1101/2022.08.18.504394

**Authors:** Prasanth Manohar, Belinda Loh, Ramesh Nachimuthu, Sebastian Leptihn

## Abstract

Phage-antibiotic combinations to treat bacterial infections are gaining increased attention due to the synergistic effects often observed when applying both components together. This synergy has also been observed for bacteria embedded in biofilms as many phages are capable of degrading the heterogeneous material, often preventing antibiotic compounds from reaching the embedded bacteria. Most studies however focus on a single pathogen, although in many clinical cases multiple species are present at the site of infection. The aim of this study was to investigate the anti-biofilm activity of phage-antibiotic/antifungal combinations on single- and dual-species biofilms formed by the bacterium *P. aeruginosa* and the fungal pathogen *Candida albicans*, two microorganisms commonly found together in cystic fibrosis patients. The *Pseudomonas* phage Motto in combination with ciprofloxacin had significant anti-biofilm activity in disintegrating 24-hour-old pseudomonal biofilms. Also, other antibiotic combinations, such as cefotaxime, gentamicin, meropenem and tetracycline showed substantial effects on biofilms. We then compared biofilms formed by *P. aeruginosa* alone with the dual-species biofilms formed by bacteria and *C. albicans*. Here, we found that the phage together with the antifungal fluconazole was active against 6-hour-old dual-species biofilms but showed only negligible activity against 24-hour-old biofilms. Similarly, the combination of antibacterial compounds together with the phage showed no synergistic effects on biofilms formed by *P. aeruginosa* and *C. albicans*. This study lays the first foundation for potential therapeutic approaches to treat infections caused by bacteria and fungi using phage-antibiotic combinations.

## 1. Introduction

Phage therapy makes use of bacteriophages to treat bacterial infections, in particular those that cannot be treated with conventional antibiotics due to multi-drug resistance (1,2). Although most of the therapeutic applications of phages are in monotherapy or as phage cocktails, recently, the combination of phage-antibiotic is getting attention through successful *in vitro* studies (3,4). In most cases, phages and antibiotics were proven to act in synergy against bacteria (5-7). Recent investigations showed that antibiotic resensitization can occur during phage exposure in antibiotic-resisting bacterial cells (8,9).

*P. aeruginosa* is a Gram-negative pathogen known to cause mild, but also life-threatening infections of the blood, the lungs and other parts of the body, here in particular after surgery. While many of the infections are treatable, an increasing number of antibiotic-resistant *P. aeruginosa* strains are being observed. This, together with the pathogen’s ability to form biofilms can make the clinical control of infections challenging (10). Recent studies showed the efficacy of *Pseudomonas* phages as anti-biofilm agents, while the use of phage-antibiotic combinations is gaining attention due to synergistic effects (11-14). Another major challenge with *P. aeruginosa* biofilms is that they often form dual-species or multi-species biofilms (13,15).

Microbe-microbe interactions can influence various infection parameters including virulence and biofilm formation. Dual-species biofilms formed by *P. aeruginosa* and *Candida albicans* are frequently found in cystic fibrosis patients but also in patients with catheters (15). *C. albicans* alone can cause opportunistic infections and is mostly found in the gastrointestinal tract and mouth (16). Most of the bacterial-fungal dual-species biofilm infections are caused by the coexistence of *Pseudomonas*-*Candida* or *Staphylococcus*-*Candida* (15,16). To date, antibiotics are the physicians’ choice for the treatment against dual-species biofilm infections but the recently increasing rates of antibiotic resistance can make the therapeutic approach unsuccessful.

To understand the potential of using phage-antibiotic combinations on *Pseudomonas*-*Candida* embedded in biofilms, we tested various antibiotics and fungicides on single and dual-species biofilms. First, we systematically tested the effects of phages together with antibiotics on planktonic, then on biofilm-embedded bacteria before we studied the impact of antimicrobials in conjunction with phages on *Pseudomonas*-*Candida* biofilms.

## 2. Materials and methods

### 2.1 Bacteria, Candida and bacteriophages

In this study, pathogenic mucoid *P. aeruginosa* PA01 and *C. albicans* C11 were used which were previously collected by the diagnostic centres in Chennai, Tamil Nadu, India. Minimal inhibitory concentration (MIC) tests were performed using cefotaxime, ciprofloxacin, gentamicin, meropenem and tetracycline (Merck KGaA, Darmstadt, Germany) (17). The results were interpreted according to CLSI guidelines (18). Previously characterized lytic *Pseudomonas* phage Motto was used in this study (NCBI accession number ON843697).

### 2.2 Phage-antibiotic combination against planktonic P. aeruginosa

To study synergistic effects, the antibiotics were added at the sub-inhibitory concentrations (1/4^th^ MIC) and the phages were used at concentrations of 10^3^, 10^6^ and 10^12^ PFU/mL (6). Briefly, antibiotics were diluted to the sub-inhibitory concentrations (1/4^th^ MIC) which were mixed with the 50 µL of phages at respective concentrations in the micro-titre plates. Then, 5 µL of *P. aeruginosa* at 5×10^5^ CFU/mL was added. The plates were incubated at 37°C for 18 hours and 100 µL was removed, serially diluted (up to 10^−6^) and plated on LB agar plates. Colony-forming units (CFU/mL) were determined and compared to the control (phage and antibiotic alone). All the data are presented as mean ± standard deviation (SD) of at least three independent experiments.

### 2.3 Phage-antibiotic combination against biofilm-forming P. aeruginosa

For the anti-biofilm studies, cefotaxime, ciprofloxacin, gentamicin, meropenem and tetracycline were used. Briefly, biofilms were formed using the exponentially growing bacterial cells from the overnight culture. Overnight cultures were grown at 37°C in LB broth and diluted in a microtiter plate and incubated at 37°C for 24 hours. The phage-antibiotic combinations were prepared following the checkerboard procedure (antibiotics from 0.5 to 32 µg/mL were prepared in columns and the phage concentrations from 10^2^ to 10^9^ PFU/mL were prepared in rows). Then, the plate was incubated at 37°C for 24 hours, washed twice to remove the planktonic cells and stained using crystal violet. The optical density (OD) was determined at 595 nm using a microtitre plate reader (BioTek, India).

### 2.4 Phage-antibiotic effects on dual-species biofilms

To study the inhibitory effect of *Pseudomonas* phage Motto and antimicrobial compounds including the fungicide fluconazole against *P. aeruginosa*-*C. albicans* co-infection, we allowed a dual-species biofilm to be formed: Briefly, overnight cultures of *P. aeruginosa* PA01 were grown at 37°C in LB broth, and overnight cultures of *C. albicans* C11 were grown at 30°C with shaking in YPD (Yeast extract, Peptone, and Dextrose) medium. For dual-species biofilms, equal volumes (∼10^5^ CFU) of *P. aeruginosa* and *C. albicans* were mixed and incubated at 37°C for 6 and 24 hours. Plates were then washed twice to remove the planktonic cells. Briefly, phages at 10^2^ to 10^12^ PFU/mL and fluconazole from 2 to 128 µg/mL were added (fluconazole in columns and phages in rows) and transferred to dual-species biofilm plates either after 6 hours or 24 hours. After the addition of phage-fluconazole combinations, the plates were incubated for 16 hours, washed, and stained using crystal violet and OD_595nm_ was measured. The control groups include pseudomonal biofilm without challenge, *C. albicans* biofilm without challenge, and dual-species biofilm challenged with phage alone and fluconazole alone. All the data are presented as mean ± standard deviation (SD) of at least three independent experiments.

In the case of phage-antibacterial agents, the dual-species biofilms (24 hours old) were treated with phages at 10^2^ to 10^9^ PFU/mL and antibacterials (cefotaxime, ciprofloxacin, gentamicin, meropenem and tetracycline) at 0.5 to 128 µg/mL. Accordingly, the dual-species biofilms (24 hours old) were treated with fluconazole at 0.5 to 64 µg/mL and antibacterials (cefotaxime, ciprofloxacin, gentamicin, meropenem and tetracycline) at 0.5 to 128 µg/mL. After the addition of phage/fluconazole-antibacterial combinations, the plates were incubated for 16 hours, washed, and stained using crystal violet and OD_595nm_ was measured.

## 3. Results

### 3.1 Antibiotics act synergistically with phage on planktonic cells

Phage-antibiotic combinations have attracted attention in the treatment of multidrug-resistant bacterial infections due to often observed synergistic effects *in vitro* and *in vivo*. We thus aimed to test the effect of phage Motto on its host in the presence of antibiotics. First, we determined the MIC of *P. aeruginosa* PA01 against different antibiotics. Five antibiotics such as cefotaxime, ciprofloxacin, gentamicin, meropenem and tetracycline were used in this study, which is either front-line or broad-spectrum antibiotics against *P. aeruginosa* infections. The isolate was found to be resistant to cefotaxime, gentamicin, meropenem and tetracycline but susceptible to ciprofloxacin according to the CLSI guidelines (Table 1). For the following experiments, sublethal concentrations of antibiotics were chosen and accordingly, 1/4^th^ of the MIC was selected. Next, all the antibiotics were tested for their phage inhibitory effects to ensure that the phage activity was not affected by the compounds. The effect of *Pseudomonas* phage Motto and sublethal concentrations of the antibiotics on *P. aeruginosa* PA01 was then investigated by determining the viable cell concentration (CFU/mL) after 18 hours of exposure. In the control group (without phage and antibiotic), the bacterial count was 3.4 × 10^9^ CFU/mL, where the phages with the final titer of 10^3^ and 10^6^ PFU/mL reduced the bacterial count to 2.1 × 10^6^ CFU/mL and 1.2 × 10^3^ CFU/mL, respectively (Fig.1). No bacteria were found when the pathogen was exposed to phages at a titre of 10^12^ PFU/mL. In the absence of phages, antibiotics at sublethal concentrations (1/4^th^ of the MIC) had minor or even no inhibitory effects. Notable bacterial reductions were observed in the case of tetracycline (4.1 × 10^7^ CFU/mL), cefotaxime (3.7 × 10^8^ CFU/mL) and gentamicin (2.1 × 10^8^ CFU/mL) but no effect was found when ciprofloxacin was used (1.1 × 10^9^ CFU/mL). Likewise, meropenem had no significant effect (2.9 × 10^9^ CFU/mL). While we observed synergistic effects of phages with all antibiotics, interestingly, we found that the combination of phage with ciprofloxacin and meropenem showed a higher degree of synergy compared to the other antibiotics we tested. The combination of ciprofloxacin (0.5 µg/mL) together with phages at 10^3^ PFU/mL resulted in a seven-log reduction. Similarly, meropenem (8 µg/mL) together with phages at 10^3^ PFU/mL resulted in a six-log reduction. Phages at a higher titre of 10^6^ PFU/mL (plus ciprofloxacin or meropenem) resulted in the complete elimination of bacterial cells. Cefotaxime-phage had synergistic activity resulting in a five-log (10^3^ PFU/mL) and six-log (10^6^ PFU/mL) reduction. The combination of 10^3^ PFU/mL phages together with gentamicin (4 µg/mL) resulted in a four-log reduction, and at 10^6^ PFU/mL a six-log reduction was observed. Tetracycline-phage combinations substantially reduced the number of viable cells: At a phage number of 10^3^ PFU/mL a four-log reduction was seen, while a higher titre of 10^6^ PFU/mL resulted in a five-log reduction of bacterial cells. As noted previously, under all the conditions no viable bacterial growth was observed at 10^12^ PFU/mL.

**Table 1:** Representation of the minimal inhibitory concentration (MIC) results of *P. aeruginosa* PA01 strain and the sub-inhibitory concentrations chosen for the study.

| S.no | Antibiotics | MIC (µg/mL) | 1/4<sup>th</sup> sub-inhibitory MIC (µg/mL) |
| --- | --- | --- | --- |
| 1. | Cefotaxime | **64** | 16 |
| 2. | Ciprofloxacin | 2 | 0.5 |
| 3. | Gentamicin | **16** | 4 |
| 4. | Meropenem | **32** | 8 |
| 5. | Tetracycline | **32** | 8 |

The resistance breakpoint (µg/mL) of Cefotaxime is ≥64, Ciprofloxacin is ≥4, Gentamicin is ≥16, Meropenem is ≥16, Tetracycline is ≥16 according to CLSI guidelines. The bolded numbers represent resistance.

**Figure 1:**
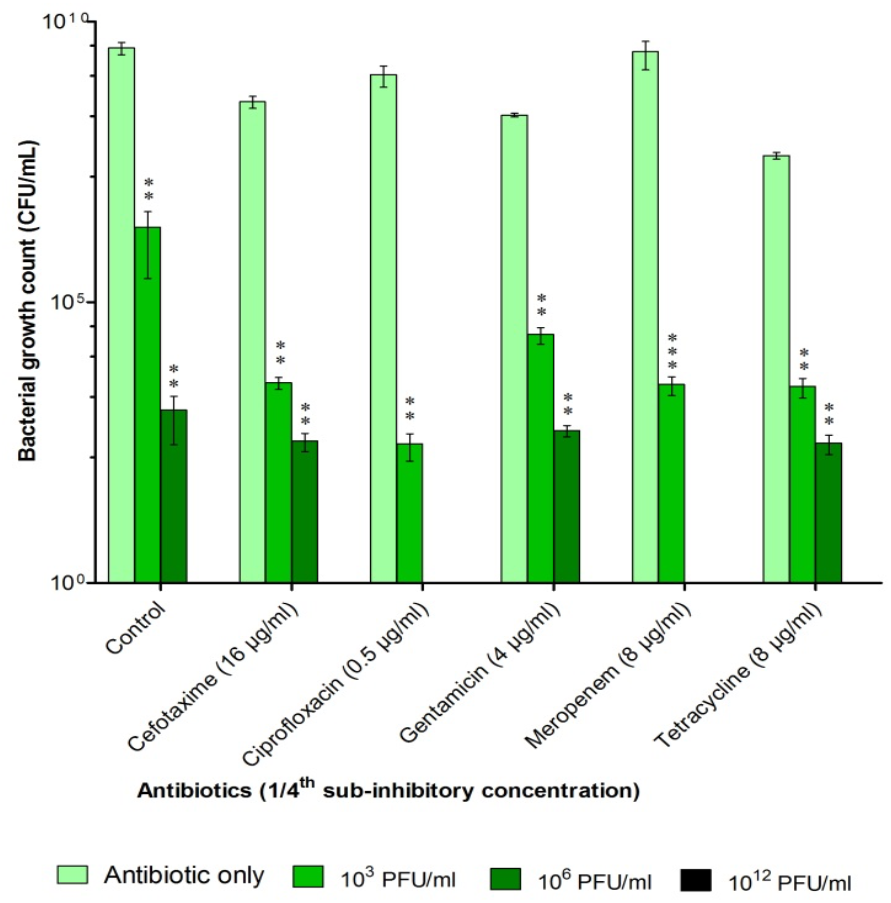
Synergistic effect of *Pseudomonas* phage Motto with antibiotics as determined by the statistical analysis. The combination provides a significant reduction in bacterial growth as observed by CFU/mL. In case no bars are visible, the bacterial count was zero. All the error bars indicated are standard errors and the ***p<0.001, **p>0.05 were determined by two-way ANOVA (between antibiotic only and phage-antibiotic combination) with Bonferroni correction.

### 3.1 Phage-antibiotic combinations show synergy against biofilm-forming P. aeruginosa cells and biofilms

Often phages and antibiotics have different effects on the same bacterial strain depending on whether they are embedded in a biofilm or planktonic. Thus, we determined the possible effects of phage-antibiotic combinations on bacterial biofilm-embedded cells. After forming biofilm-embedded cells in microtiter plates for 24 hours, we added different concentrations of antibiotics together with different amounts of phages. A synography model was previously designed by the TAILΦR lab in Texas, to study and visualise the effectiveness of phage-antibiotic combinations across many stoichiometries (4).

The analysis of our data shows that for all phage-antibiotic combinations, at high titres and antibiotic concentrations, the biofilm is reduced to a varying extent and depending on the antibiotic. In the treatment of the pseudomonal biofilm with cefotaxime alone, even the highest antibiotic concentration of 32 µg/mL did not show any significant anti-biofilm effect. However, complete removal of the biofilm was observed at concentrations of cefotaxime between 32 and 16 µg/mL when phages were present at high titres (Fig.2A). MIC results showed that the *P. aeruginosa* PA01 was sensitive to ciprofloxacin. Correlating with this, we observed a complete biofilm eradication at the higher concentrations of ciprofloxacin (>= 16 µg/mL) which appears at first glance surprising (Fig. 2B). Possibly the bactericidal effects lead to a release of bacterial enzymes from the cells which then degrade the biofilm as a direct effect of the ciprofloxacin is less likely; this interpretation is speculation and requires further experimental validation. Synergistic effects were observed with increasing amounts of phages. Gentamicin alone exhibited minor anti-biofilm properties but combinatorial effects can be seen when phages are added at titres higher than 10^3^ PFU/mL (Fig.2C). Similarly, meropenem, which had no obvious effect on biofilms on its own, did only reduce the biofilm when phages were present. Clear synergy was observed at values higher than 4 µg/mL and titres higher than 10^7^ PFU/mL (Fig.2D). While tetracycline alone showed -to some extent-anti-biofilm properties in our assay, the combination of the antibiotic with the phage exhibited significant synergistic effects at concentrations higher than 4 µg/mL and phage numbers higher than 10^7^ PFU/mL (Fig.2E).

**Figure 2:**
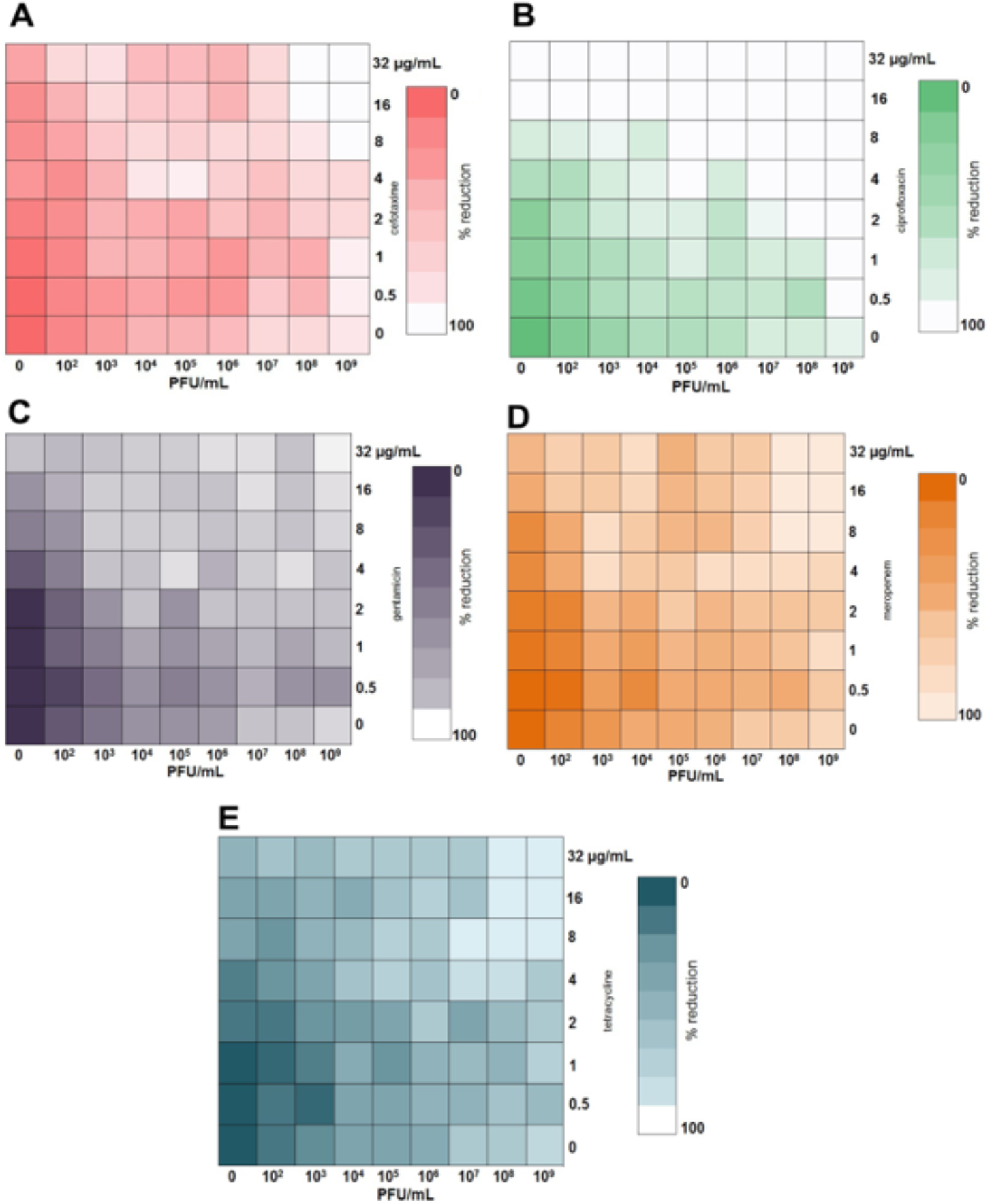
Anti-biofilm effect of *Pseudomonas* phage Motto and different antibiotics on phage-antibiotic synergy. The effect of different antibiotics was studied: (A) Cefotaxime; (B) Ciprofloxacin; (C) Gentamicin; (D) Meropenem; (E) Tetracycline. The biofilms (24 hours old) were treated with different combinations of phages and antibiotics. The synograms represent the OD_595nm_ values read after 24 hours of treatment and the mean reduction percentage of each treatment from three independent replicates.

### 3.3 Combinations of phage and the antifungal fluconazole have minor effects on dual-species biofilms

To investigate the formation of mono- and dual-culture biofilms, we cultured cells of *P. aeruginosa* PA01 and *C. albicans* C11 either alone or together in microtiter plates. After 6 hours, cultures of *P. aeruginosa* PA01 quantitatively formed less biofilm compared to *C. albicans* C11, according to the assay we employed. Both organisms together, however, formed even more biofilm than the individual ones combined (Fig.3A). After 24 hours, biofilms increased for both the bacterial and the fungal pathogen, while the dual-species biofilm was less than the sum of the individual biofilms from *P. aeruginosa* PA01 and of *C. albicans* C11 (Fig.3B).

**Figure 3:**
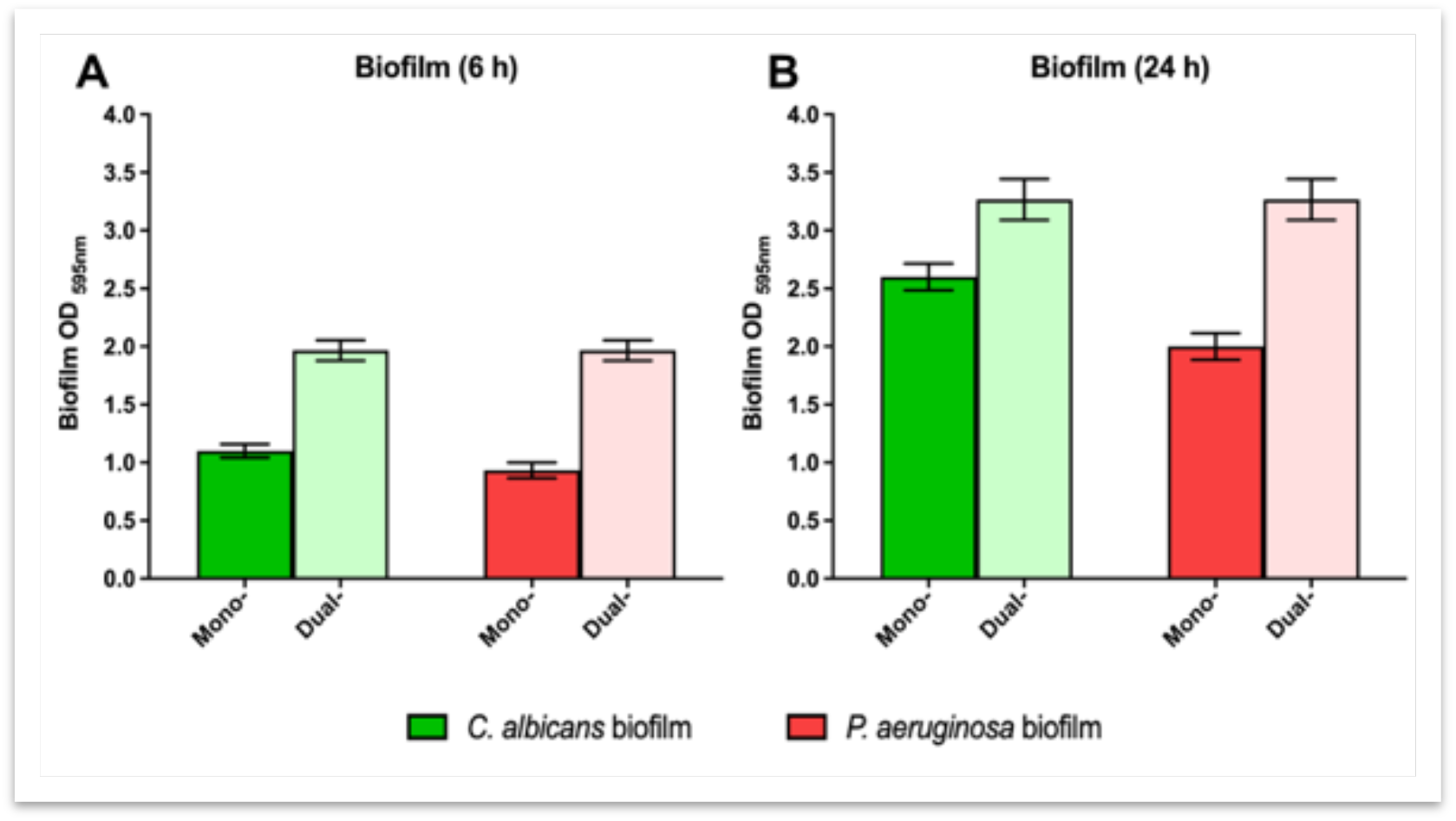
Biofilm of *C. albicans* and *P. aeruginosa* clinical isolates in mono- and dual-culture at 6 h (left) and 24 h (right). The results represent the biofilm-formation after 6 and 24 hours as the absorbance was read at OD_595nm_ and the error bars represent the standard mean of the three independent experiments.

The eradication of dual-species biofilms is impossible in the presence of phage alone or antibiotics alone as the fungal pathogen can multiply and continue to produce biofilm. Therefore, we tested phage-antifungal combinations. A complete eradication was not observed, even at the highest phage and fluconazole concentrations (Fig.4). Nonetheless, a trend at high phage and antibiotic concentrations with a biofilm-adverse effect can be observed in both cases of up to 30%, with 6 and 24-hour biofilm samples (Fig.4A, B). It is clear that the phage has a positive impact on the removal of the dual-species biofilms in combination with the exposure to fluconazole.

**Figure 4:**
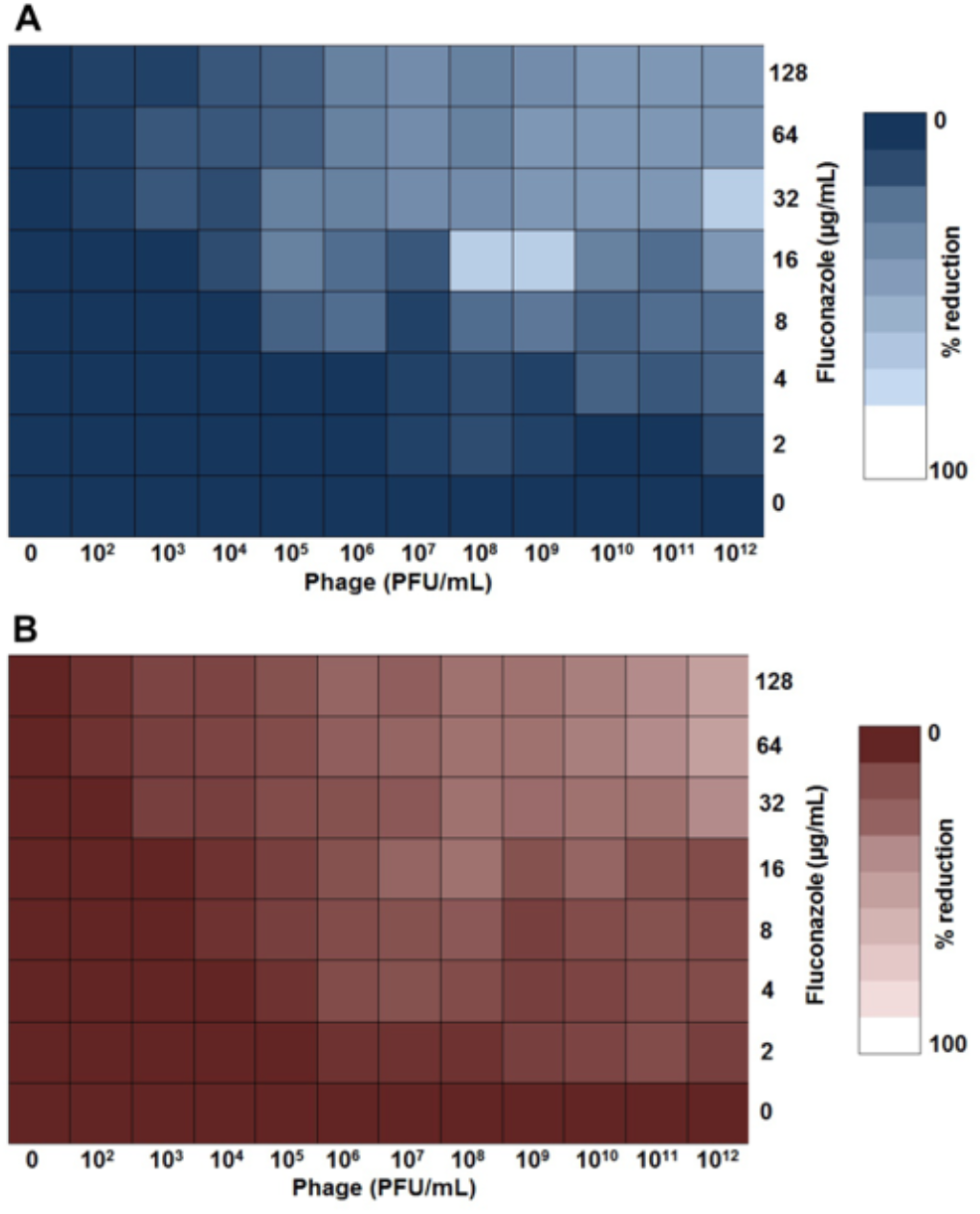
Phage-fluconazole synergism in the reduction of dual-species biofilms formed by *Pseudomonas*-*Candida*. Effect of *Pseudomonas* phage Motto (10^2^ to 10^12^) and fluconazole (128 to 2 µg/mL) on dual-species biofilms. The dual-species biofilms [6 hours old (top) and 24 hours old (bottom)] were treated with different combinations of *Pseudomonas* phage and fluconazole. The synograms represent the OD_595nm_ values as read after 24 hours of treatment and the mean reduction percentage of treatment from three independent replicates.

### 3.4 Antifungals are required for dual-species biofilm removal, as phage-antibacterial combinations have no effect

We established that phage-antibiotic combinations are effective on biofilms formed by *P. aeruginosa* alone. We also demonstrated that phage-antifungal combinations resulted in the reduction of biofilms formed by the bacterial pathogen together with *C. albicans*, albeit to a minor extent. Thus, we next tested if the combination of phages together with antibiotics acting on *P. aeruginosa* leads to a reduction of dual-species biofilms. However, even at the highest concentration and highest phage titre tested, biofilms remained unaltered (Fig.5). This is somewhat surprising, as one might suspect that the bacterial biofilm is reduced. However, possibly the fungal pathogen continues to thrive and thus produces more biofilm.

**Figure 5:**
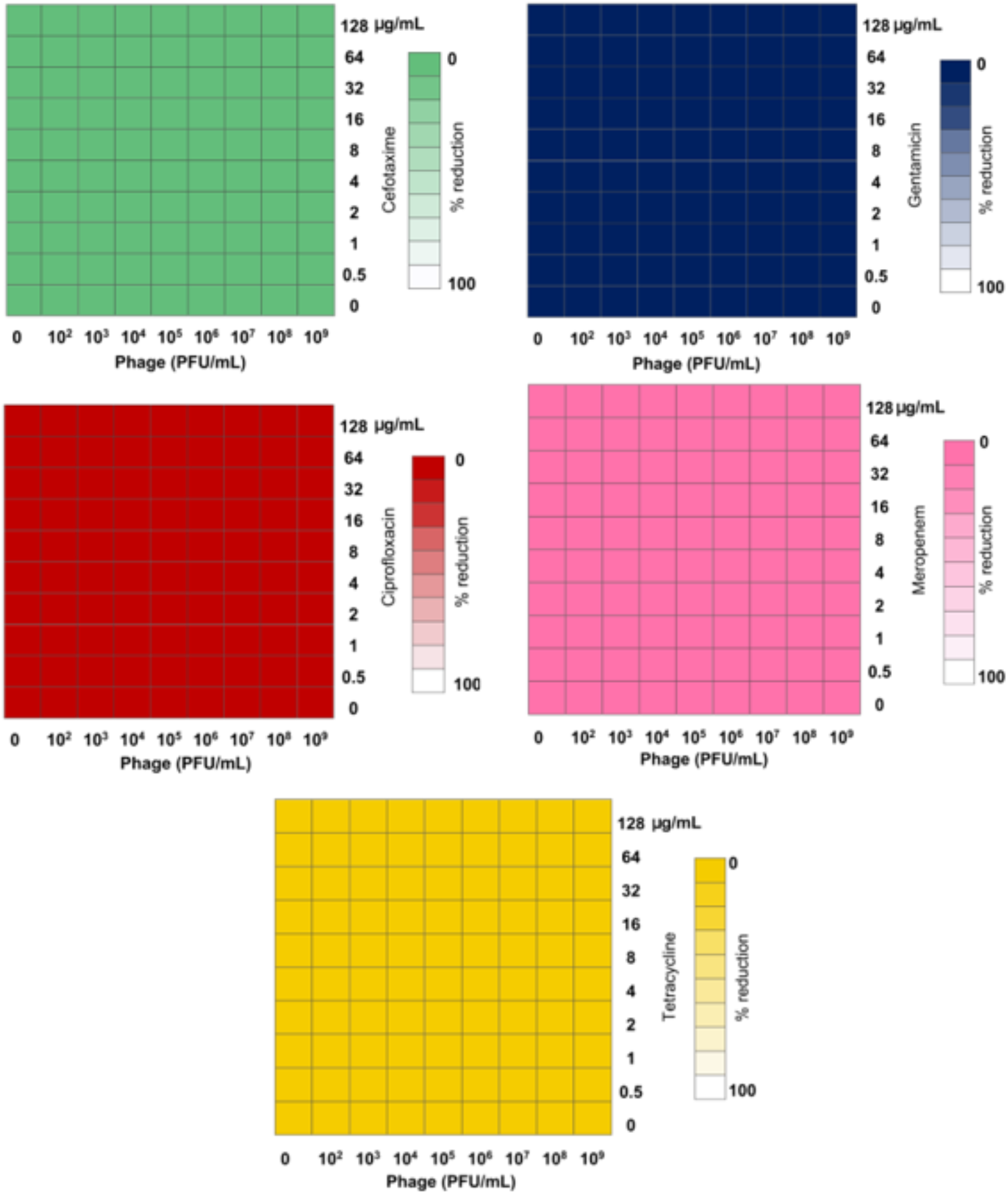
Phage-antibiotic synergism in the reduction of dual-species biofilms formed by *Pseudomonas*-*Candida*. Effect of *Pseudomonas* phage Motto (10^2^ to 10^9^) and antibiotics (cefotaxime, ciprofloxacin, gentamicin, meropenem, tetracycline) on dual-species biofilms. The dual-species biofilms (24 hours old) were treated with different combinations of *Pseudomonas* phage and antibiotics. The synograms represent the OD_595nm_ values as read after 24 hours of treatment and the mean reduction percentage of treatment from three independent replicates.

### 3.5 Antimicrobial combinations of the antifungal fluconazole together with antibacterials act synergistically at high concentrations

Although this study focused on the antibiofilm properties of a *P. aeruginosa* phage in combinations with antimicrobial compounds, we also investigated how antibiotics alone act on dual-species biofilms. Interestingly, we observe antibiotic-specific synergy in some cases; clear effects can be seen in the case of cefotaxime and ciprofloxacin, while the reduction of biofilm using gentamicin together with fluconazole was less pronounced. The effect with meropenem in conjunction with the antifungal was even more obvious, while the fluconazole-tetracycline combination was unable to reduce the biofilm altogether (Fig.6).

**Figure 6:**
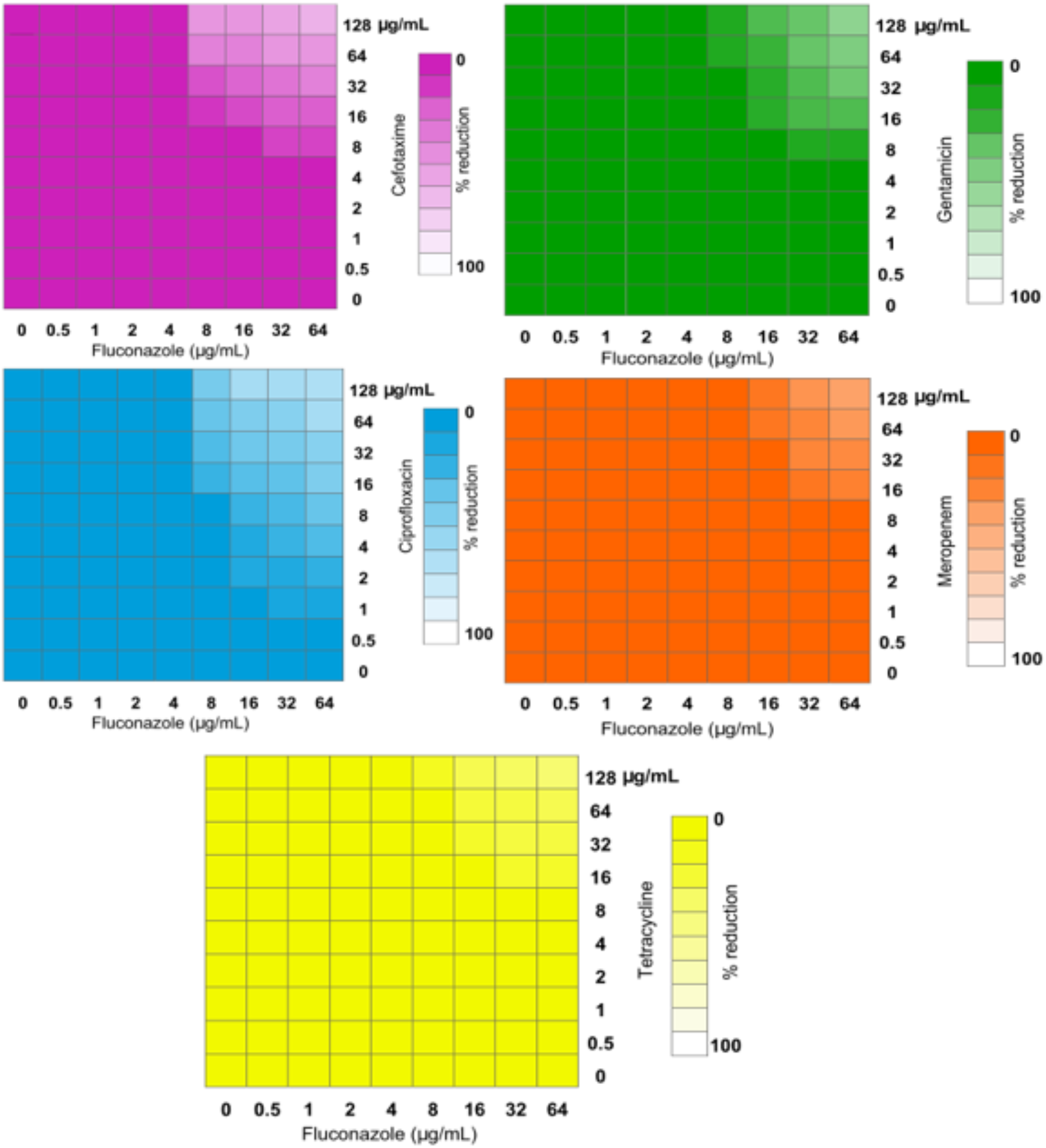
Fluconazole-antibiotic synergism in the reduction of dual-species biofilms formed by *Pseudomonas*-*Candida*. Effect of fluconazole (0.5 to 64 µg/mL) and antibiotics (cefotaxime, ciprofloxacin, gentamicin, meropenem, tetracycline at 0.5 to 128 µg/mL) on dual-species biofilms. The dual-species biofilms (24 hours old) were treated with different combinations of fluconazole and antibiotics. The synograms represent the OD_595nm_ values as read after 24 hours of treatment and the mean reduction percentage of treatment from three independent replicates.

## 4. Discussion

*P. aeruginosa* is known to cause life-threatening infections in humans and is one of the common nosocomial pathogens to cause respiratory tract infections. To overcome the threat of antibiotic-resistant pseudomonal infections, non-antibiotic therapies and antibiotic combinatorial therapies are being investigated as valuable alternatives (19). The use of phage-antibiotic combination therapies has been therapeutically successful, possibly due to the emergence of phage-resistant strains that show increased antibiotic susceptibility (7,8,20). However, many infections are complicated by the formation of biofilms, which inhibit the diffusion of antibiotic compounds and form a kind of protective layer around the pathogens. Due to their unique properties, phages have been found to be effective in dissolving biofilms, both *in vitro* and *in vivo* (21,22). This was also demonstrated in the case of *P. aeruginosa* infections (23,24). Thus, phages can act synergistically as a biofilm-disrupting agent, allowing antibiotics to penetrate the target. Despite this, many infections are caused not by one species but by several, known as polymicrobial infections. Such “collaborative infections” are common for *P. aeruginosa* which often is found together with *S. aureus* (25). Especially in the lungs of cystic fibrosis (CF) patients, *P. aeruginosa* is present with *C. albicans*. Thus, there is growing interest in multispecies biofilms. This is particularly important for those that are formed from different domains of life. The treatment failures are mainly due to the impermeability of dual-species biofilms by chemotherapeutic drugs.

Our study aimed to investigate the potential of phage-antibiotic combinations in eradicating biofilms formed by *P. aeruginosa* and dual-species biofilms of *P. aeruginosa*-*C. albicans*. Initially, *P. aeruginosa* PA01 planktonic cells were treated with phage-antibiotic combinations in order to establish a baseline for potentially observed phage-antibiotic synergy. Here, we found that ciprofloxacin and meropenem had a more pronounced synergistic effect. On the other side of the spectrum, cefotaxime, gentamicin and tetracycline had the least degree of synergism. Despite the bacteria being susceptible to ciprofloxacin (MIC of 0.5 µg/mL), when combined with phages even at the sublethal concentrations (1/4^th^ MIC), a lethal anti-pseudomonal effect was observed. The other antibiotics also had a synergistic effect with a minimum of four-log reductions when compared to antibiotics alone and at least two-log reductions when compared to phages alone. Notably, irrespective of the antibiotic class, the *Pseudomonas* phage Motto was found to work synergistically with all the tested antibiotics in this study. Our data is similar to a recent study which showed the anti-pseudomonal effect of phage-antibiotic (sub-MIC) combinations on pathogenic *P. aeruginosa* strains (20). As we were interested in the effects on biofilm, we tested the reduction of pseudomonal biofilms when exposed to phage-antibiotic combinations. Here, we observed that phage-ciprofloxacin exhibited enhanced biofilm eradication compared to other antibiotics (cefotaxime, gentamicin, meropenem and tetracycline). Similar results were reported previously showing that the phage-ciprofloxacin combination has better biofilm eradication ability compared to the single-compound treatment (14). In our study, the synogram showed that phage Motto was acting synergistically with all tested antibiotics and was effective at disrupting *P. aeruginosa* biofilms.

As *P. aeruginosa* is often found together with *C. albicans* in biofilms, we focussed on this important aspect next: The inhibition of *P. aeruginosa*-*C. albicans* dual-species biofilms when exposed to the combination of phage together with the fungicide fluconazole were tested. To this end, we investigated the formation of biofilm by the pathogens on their own or when growing together. Curiously, while *P. aeruginosa* PA01 and *C. albicans* C11 were forming moderate amounts of biofilm within 6 hours, we observed that the production of biofilm was additive when both pathogens were present. More substantial biofilms were produced by the individual species after 24 hours, while the dual-species biofilm reached even higher quantities in our assay, yet not the sum of the individual amounts of mono-species biofilms.

After establishing the extent of the formation of biofilms at the two-time points, we next tested the combination of the phage Motto together with the antifungal compound fluconazole. While complete biofilm eradication was observed in the case of 6-hour-old biofilms, the 24-hour-old biofilms were only partially (50%) disintegrated after 16 hours of phage-fluconazole exposure. A possible explanation for this strong difference is the change in biofilm composition over time, which remains to be investigated. While many phages are known to degrade bacterial biofilms, it is less likely that depolymerases produced by phages can degrade dual-species biofilms, especially the ones made by fungi.

We also tested the combination of antibacterial compounds together with the antifungal drug. Here, we observed a clear synergy albeit at high concentrations of both compounds. To our surprise, no effects at all were seen when phage antibacterial combinations were used to treat the dual-species biofilm. Our study aimed to shed light on the complexity of polymicrobial biofilms and the effects of phages on their structure. It becomes obvious that we need more studies to understand the effect of phages on such complex systems in order to develop treatment regimes for dual-species infections or biofilm removal before phages can be considered for therapeutic purposes.

## Acknowledgement

The authors would like to thank the Vellore Institute of Technology for the partial support provided by the VIT-SEED grant.

## Author’s contribution

PM participated in the study design, research work, preliminary data collection and data analysis and drafted the manuscript. NR and SL participated in the design of the study. NR, BL and SL performed statistical analyses and wrote the manuscript. All the authors read and approved the final manuscript.

